# Orthogonal representation of task-related information in theta phase-based multiple place fields of single units in the subiculum

**DOI:** 10.1101/2021.08.11.456028

**Authors:** Su-Min Lee, Jae-Min Seol, Inah Lee

## Abstract

The subiculum is positioned at a critical juncture at the interface of the hippocampus with the rest of the brain. However, the exact roles of the subiculum in most hippocampal-dependent memory tasks remain largely unknown. One obstacle to make analytical comparisons of neural firing patterns between the subiculum and hippocampal CA1 is the broad firing fields of the subicular cells. Here, we used spiking phases in relation to theta rhythm to parse the broad firing field of a subicular neuron into multiple subfields to find the unique functional contribution of the subiculum while male rats performed a hippocampal-dependent visual scene memory task. Some of the broad firing fields of the subicular neurons were successfully divided into multiple subfields by using the theta-phase precession cycle. The resulting phase-based fields in the subiculum were more similar to those in CA1 in terms of the field size and phase-precession strength. The new method significantly improved the detection of task-relevant information in subicular cells without affecting the information content represented by CA1 cells. Notably, multiple fields of a single subicular neuron, unlike those in the CA1, could carry heterogeneous task-related information such as visual context and choice response. Our findings suggest that the subicular cells integrate multiple task-related factors by using theta rhythm to associate environmental context with action.

## Introduction

The hippocampal formation plays key roles in fundamental cognitive functions, including spatial navigation and episodic memory (Scoville and Milner, 1957; O’Keefe and Nadel, 1978; Eichenbaum, 2000). The subiculum, a region within the hippocampal formation, has long been considered the area from which cortical outputs of the hippocampus emanate (Amaral et al., 1991; Witter et al., 2000). However, viewing the subiculum as an area that passively transmits hippocampal information to cortical regions might be inappropriate, because the subiculum is connected not only with the CA1 of the hippocampus but also with other areas, including the medial prefrontal cortex, entorhinal cortex, retrosplenial cortex, perirhinal cortex, postrhinal cortex, nucleus accumbens, basal amygdala and various subcortical regions (Witter, 2006; Cembrowski et al., 2018b; Matsumoto et al., 2019).

Physiologically, it has been reported that the neural correlates of the subiculum are significantly different from those of the CA1 during spatial navigation. Specifically, neurons in the subiculum tend to exhibit broader place fields than those in the CA1 (Barnes et al., 1990; Sharp and Green, 1994; Kim et al., 2012b; Lee et al., 2018). Also, place cells in the subiculum are more attuned to movement-related factors, such as direction and motion, during navigation compared with CA1 place cells (Lever et al., 2009; Olson et al., 2017; Kitanishi et al., 2021; Ledergerber et al., 2021). A few studies have also suggested that the subiculum is essential in remembering places and environmental contexts (Morris et al., 1990; Potvin et al., 2007, 2009; Potvin et al., 2010; Melo et al., 2020). However, the exact roles of subicular neurons, especially in a goal-directed memory task, still remain largely unknown. In our previous study, we reported that neurons in both the subiculum and CA1 showed rate remapping according to task-related factors, specifically visual scene and choice response in a visual scene memory (VSM) task in which rats were required to make choices in a T-maze using the visual scene stimulus presented around the maze (Lee et al., 2018). Interestingly, place cells in the CA1 showed such firing properties while coding very specific locations in space, whereas cells in the subiculum fired similarly while mapping broader areas (e.g., stem or choice arm region), as if they represent the cognitive structure of the task by schematically parsing the environment. On the basis of these results, we speculated that position-linked environmental information in the hippocampus in the VSM task (Delcasso et al., 2014; Lee and Lee, 2020) might be translated into contextual action-related information that can be communicated with other brain regions.

One major obstacle that poses great difficulties for investigations of the neural correlates of subicular neurons is their higher spontaneous firing rates and broader firing fields in space compared with those of place cells in the hippocampus (Sharp and Green, 1994; Kim et al., 2012b; Lee et al., 2018). These firing characteristics of subicular neurons make it difficult to apply the conventional analytical techniques optimized for place cells recorded from hippocampus, where place fields are more restricted to specific locations of the environment with a higher signal-to-noise ratio compared with the subiculum. For example, in our previous study (Lee et al., 2018), we sought to identify field boundaries of subicular cells by finding local minima through statistical comparisons of trial-by-trial firing rates between neighboring bins. However, such methods had shortcomings, such as defining some subicular cells as having no fields and ignoring small subfields in the presence of a dominant field with a very high firing peak.

Notably, some previous studies attempted to parse the broad spatial firing field of a subicular neuron into smaller fields using the phases of spikes in relation to theta rhythm (Maurer et al., 2006; Kim et al., 2012b). Here, inspired by these studies, we compared the traditional rate-based field-detection method with the theta phase-based field-detection method using the same physiological data recorded from the CA1 and subiculum in our previous study (Lee et al., 2018). The current study showed that the phase-based analysis could successfully parse subicular firing fields into multiple subfields and that these newly parsed place fields in the subiculum better represented task-related information. Importantly, some subicular cells represent multiplex information associated with the VSM task through their phase-based subfields, possibly suggesting a unique role of the subiculum in integrating environmental information with action.

## Materials and Methods

### Subjects

Male Long-Evans rats (n = 5) were used in the current study. Food was restricted to maintain rats’ body weights at 350–400 g (85% of free-feeding weight), and water was made available *ad libitum*. Rats were individually housed under a 12-h light/dark cycle. All protocols were approved by the Institutional Animal Care and Use Committee of Seoul National University.

### Behavioral task

Detailed descriptions of our experimental procedures, including the visual scene memory (VSM) task and the apparatus (**Figure 1A**), are available in our previous study (Lee et al., 2018). Briefly, the rat was located in a start box before a trial began. The experimenter started the trial by opening the door of the start box, which also triggered presentation of a patterned visual stimulus (i.e., visual scene) in an array of three adjacent LCD monitors surrounding the choice-arm region of the T-maze. The rat then entered and ran along the stem of the T-maze (stem, 73 × 8 cm; arms, 38 × 8 cm) and was required to turn left or right at the end of the stem (*choice point*) in association with the visual scene. The rat obtained a quarter piece of cereal reward (Froot Loops, Kellogg’s) from the food well at the end of the correct arm, but no reward was given if it entered the wrong arm. Four visual scenes (zebra, bamboo, pebbles, mountains) were used. In all sessions, zebra stripes and bamboo patterns were associated with the left arm, and pebbles and mountain patterns were associated with the right arm; within a session, the four visual scenes were presented in a pseudorandom sequence.

**Figure 1.**
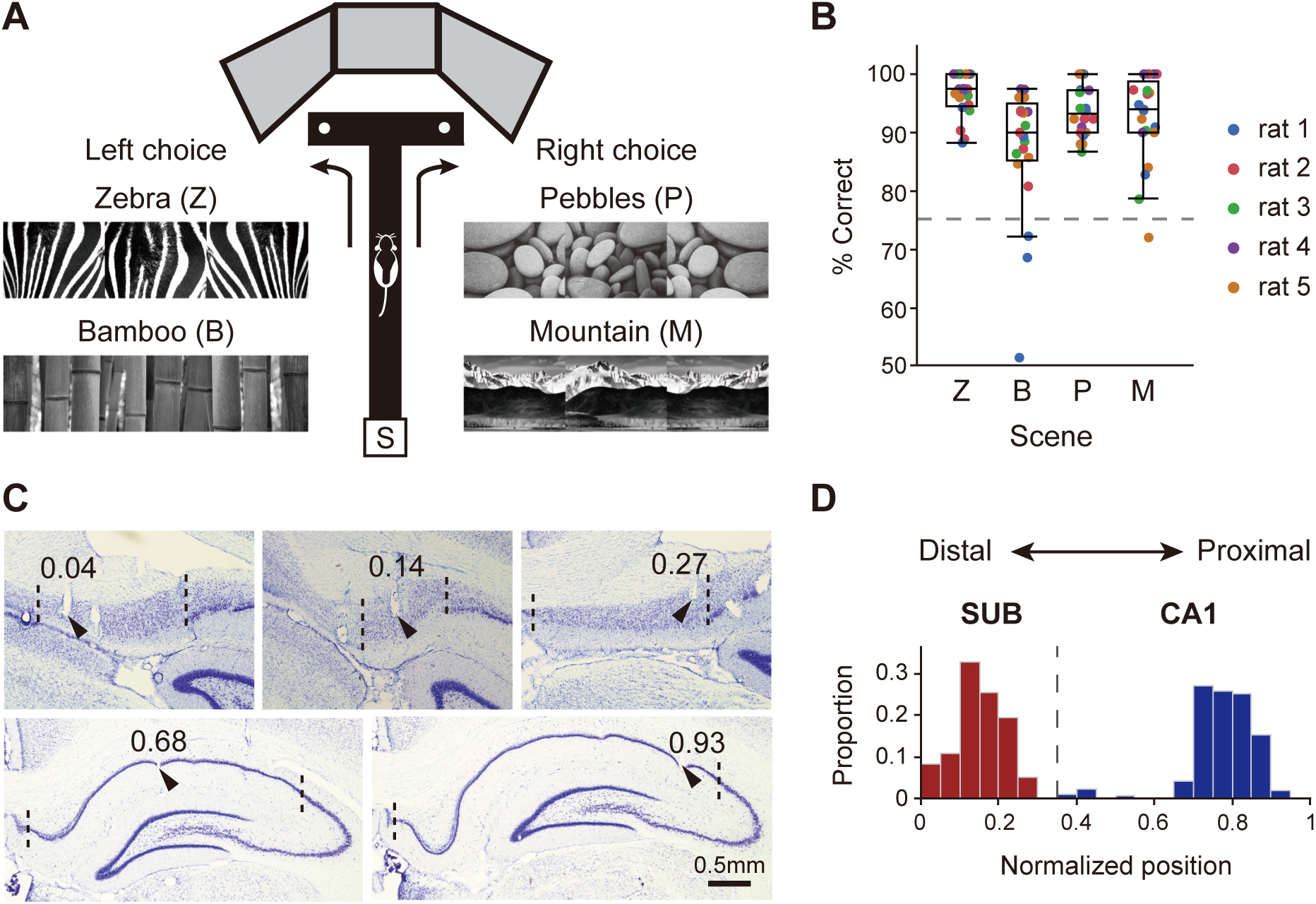
Behavioral task and histological verification of electrophysiological recordings. (**A**) VSM task. As a trial begins, the rat runs out onto the track of a T-maze from the start box (S), and one of four visual scene stimuli (Zebra, Z; Bamboo, B; Pebbles, P; Mountain, M) is presented on LCD monitors. Each scene stimulus is associated with either the left or right arm of the T-maze. The rat obtains a piece of cereal reward from the food well, located at the end of both arms, if it chooses the correctly associated side. (**B**) Behavioral performance during recording sessions (21 sessions from 5 rats). Each dot corresponds to the percent correct for each scene stimulus of a session and is color-coded for individual rats. Box plot indicates interquartile range and median value. The median values exceeded the performance criterion (dashed line, 75%) for all scenes. (**C**) Photomicrographs of Nissl-stained coronal brain sections with verified electrode tips (black arrows). Numbers above the arrows indicate normalized positions of marked recording sites along the proximodistal axis. Dashed lines represent the anatomical boundaries of the CA1 and subiculum. Upper and lower rows show recording sites from the subiculum and CA1, respectively. (**D**) Proportional distribution of cells recorded in the CA1 (blue) and subiculum (SUB; red) along the proximodistal axis (CA1, n = 270; SUB, n = 151). The positions are normalized to account for differences in relative length between two regions (see **Methods)**. The dashed line at 0.36 indicates the boundary between two regions.

During the pre-surgical training period, the rat was initially trained with a pair of scene stimuli (zebra vs. pebbles or bamboo vs. mountain, counterbalanced for rats) until it reached the performance criterion for each pair (≥ 75% correct for each scene for two consecutive days). Once the rat reached the performance criterion for both scene pairs, a hyperdrive carrying twenty-four tetrodes (+3 reference electrodes) was surgically implanted in the right hemisphere to cover 3.2-6.6 mm posterior to bregma and 1-4 mm lateral to the midline. After 1 wk of recovery, the rat was retrained until it reached pre-surgical performance levels (**Figure 1B**), during which time the tetrodes were lowered into the subiculum and CA1 by 40–160 μm daily. Thereafter, the main recording sessions (123 ± 6 trials/session, mean ± SEM) began, and the four scene stimuli were presented in an intermixed fashion during sessions.

### Electrophysiological recording and histological procedures

Single unit spiking activity and local field potentials (LFPs) were recorded from the dorsal CA1 and subiculum. Neural signals were transmitted to the data acquisition system (Digital Lynx SX; Neuralynx) through a headstage connected to the EIB board and tethered via a slip-ring commutator on the ceiling. Neural signals from tetrodes were amplified 1,000–10,000 times and sampled at 32 kHz. Spiking data were acquired by filtering at 600–6,000 Hz. LFPs were obtained by filtering the same signals at 0.1–1,000 Hz. After completion of all recording sessions, electrolytic lesions (10 μA current for 10 s) were made to mark the tip positions of the tetrodes. Twenty-four hours after electrolytic lesioning, the rat was sacrificed by inhalation of an overdose of CO_2_ and perfused transcardially. Brain tissue was stained using thionin or Timm’s method for Nissl substances (see details in Lee et al. (2018); **Figure 1C**).

The anatomical boundaries of the CA1 and subiculum were determined based on the rat brain atlas (Paxinos and Watson, 2009). Tetrodes located in the transition area between the CA1 and subiculum were excluded. To quantitatively describe the proximodistal positions of the recording tetrodes, we measured the linearized length of the cell layer in the CA1 and subiculum—specifically, the distance between the most distal to the proximal end along the curved pyramidal cell layer in a given section—using image processing software (ImageJ; NIH). Recording positions across rats were normalized by selecting a median value among the linearized lengths of the pyramidal cell layers of the CA1 and subiculum in all rats, and the ratio between the CA1 and subiculum was obtained (subiculum:CA1 = 0.36:0.64). The relative positions of tetrode tips within each region were then calculated (**Figure 1D**).

### Extraction of outbound running epochs

Before proceeding with a set of analyses based on spiking data in relation to their theta phases, we extracted only those epochs associated with outbound journeys (from the start box to either left or right food well). To facilitate theta rhythm-related analyses, we calculated the instantaneous running speed so as to include epochs in which rats ran at a reasonable speed. To this end, we interpolated linearized position data to compensate for vacancies caused by rat head movements and/or tether interference. Next, outlier data points were suppressed using a locally weighted robust regression. Then, the instantaneous running speed, calculated by dividing the length of three consecutive data points by the duration of time, was assigned to the middle point of the three. The average running speed was 35.3 cm/s in all sessions for all rats. Spikes that occurred when running speed was greater than 20 cm/s were used in this study. If the latency from the start box to the food well was longer than 6 s, that trial was discarded.

### Spiking data analysis

#### Unit isolation

Single units were isolated manually using both commercial software (SpikeSort3D; Neuralynx) and a custom-written program (WinClust) based on the waveform parameters, peak amplitude, energy, and peak-to-valley latencies. The same criteria from our previous study (Lee et al., 2018) were used to evaluate unit-isolation quality, with the additional criterion that the number of spikes during running epochs of outbound journeys on the track should be greater than 50 (subiculum, n = 208 units; CA1, n = 327 units).

#### Detection of firing rate-based place fields

Position data acquired during outbound running epochs were first linearized by scaling down using 2-cm spatial bins. The choice point—that is, the point where the rat’s position data diverged between left and right choice trials—was determined by detecting the spatial bin with a statistical difference between left and right position traces (two-sample t-test). Then, a linearized firing rate map was constructed by dividing the number of spikes by the number of position data points in individual spatial bins. Boundaries of a firing field were defined as the first spatial bin at which the firing rate dropped below 33% of the peak firing rate for two consecutive bins. If a local peak exceeding 50% of the maximum peak firing rate was found outside the predetermined firing field, it was considered as the peak of a possible subfield, and the boundaries of the subfield were found using the same algorithm. After defining the field boundaries, a firing field was identified as a place field when the peak firing rate within the field exceed 1 Hz and the spatial information score of the field was greater than 0.5. The spatial information score was computed according to the definition of Skaggs et al. (1993) as follows:

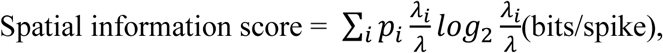

where *i* denotes the spatial bin, *p_i_* is the occupancy rate in the *i*^th^ bin, *λ_i_* is the mean firing rate in the *i*^th^ bin, and *λ* is the overall mean firing rate. The mean firing rate and peak firing rate were obtained from the raw rate maps. For display, the rate maps were smoothed using the adaptive binning method.

### LFP analysis

#### Tetrode selection

To align baseline offsets, we down-sampled LFPs from 32 kHz to 2 kHz and filtered them at 3–300 Hz using a zero-phase bandpass filter (3^rd^-order Butterworth filter with the *filtfilt* function in MATLAB). LFP traces from running epochs were then visually inspected to exclude tetrodes whose voltage traces exceeded the maximum value (3,000 μV) of the analog-digital converter or artifacts such as bumping noises. Spiking phases in relation to theta rhythm were analyzed by obtaining a power spectral density (PSD) function using a multi-taper method (Chronux ToolBox; MATLAB) and then selecting reference tetrodes with the strongest power in the high theta band (7-12 Hz) for individual sessions and regions. The frequency range of the theta band was set so as to include the most prominent peak at 8 Hz in the mean power spectral density function during the outbound journey and to minimize bumping noises that usually occurred at less than 7 Hz. LFPs recorded from the CA1 and subiculum were used for spiking phase analyses of single units in the corresponding regions.

#### Spiking theta phases

LFPs from reference tetrodes were filtered in the theta range (7-12 Hz) using a zero-phase bandpass filter, followed by application of a Hilbert transform to decompose filtered LFPs into amplitude and phase information. Spiking-phase relationships were examined by plotting instantaneous theta phases and rat’s linearized positions at time points when spikes occurred in a 2-dimensional space (phase-position plot).

#### Identification of theta phase-based place fields using DBSCAN

To define a cluster of spikes that shared the same spiking-phase relationships, we adopted a well-known clustering algorithm called the *Density-Based Spatial Clustering of Applications with Noise (DBSCAN)* suggested by Ester et al. (1996). DBSCAN is a density-based, nonparametric algorithm that gathers data points in close proximity while excluding distant or sparsely located points as noise. In DBSCAN, it is not necessary to specify the number of clusters in advance. Still, some parameters must be predetermined to run the algorithm, such as the distance (ε) and the minimum number of points within a distance (N_min_). Specifically, if the number of data points at a distance ε from a point is greater than N_min,_ including itself, the point is defined as a core point of a cluster. If another point contains the core point within distance ε but does not satisfy N_min_, it is defined as a border point. If there is no core point and N_min_ is not satisfied, the point is defined as a noise point. In our study, clusters on the position-phase plot were captured by the DBSCAN algorithm to find theta phase-based place fields (*TP-based place fields*). To avoid detecting spurious sparse clusters, we restricted DBSCAN parameters to the following ranges: distance (ε) < 8 cm; N_min_ ≥ 10; and total number of spikes in a cluster ≥ 30. The parameters were determined manually in those ranges so that the number of clusters in a cell was greater than the number of local maxima in linearized firing rate maps. Biased clustering caused by experimenter subjectivity was prevented by performing cross-validation with three additional experimenters who did not participate in analyses of the current data sets. After cross-validation, cells with invalid clustering were excluded from the analysis based on the following: (1) DBSCAN parameters not satisfied (subiculum, n = 21; CA1, n = 24), (2) insufficient spikes (subiculum, n = 17; CA1, n = 30), or (3) irregular cluster shape (subiculum, n = 16; CA1, n = 6).

#### Quantification of theta phase precession

After identification of individual place fields using theta phases, the slopes of theta precession were measured by fitting individual spike clusters to circular-linear regression lines (Kempter et al., 2012). A circular-linear correlation was also applied to determine if the phase shift was significant (Toolbox for circular statistics with MATLAB; Berens, 2009). Theta-phase precession of a place field was considered significant if the following criteria were met: (1) range of phase shift ≥ 180°, (2) slope of regression line < 0, and (3) p-value of circular-linear correlation ≤ 0.05.

### Analysis of rate remapping

To measure the amount of rate modulation between firing rate maps associated with different trial conditions (i.e., scene stimulus or choice response), we obtained a rate difference index (RDI) by calculating an absolute value of *Cohen’s d*:

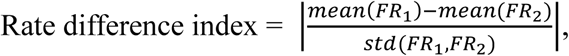

where FR_1_ and FR_2_ denote the in-field firing rates of individual trials associated with different conditions. *Cohen’s d* measure was adopted because it includes a term for standard deviation in its denominator, which was expected to control for the confounding effect induced by the variability in intrinsic firing between the CA1 and subiculum. With respect to RDI for scene stimuli, two RDI values were obtained from two pairs of scenes associated with the left or right choice arm (RDI_SCN-L_ and RDI_SCN-R_, respectively), then a maximum value was chosen as a representative scene-based RDI of a cell (RDI_SCN_). RDI for choice response (RDI_CHC_) was measured by calculating the difference in firing rates between left and right choice trials. For calculation of RDI_CHC_, only firing rate maps based on the areas ahead of the choice point were used because the rat’s actual positions on the maze were different after the choice point. Since RDI_SCN_ was originally calculated from firing rate maps associated with the same choice arm, all spiking activities after the choice point were used. However, if a spatial bin with a firing rate less than 75% of the peak firing rate was located in one arm of the maze, the field was considered an arm field and was excluded from the RDI analysis. For joint comparisons of changes in RDI_SCN_ and RDI_CHC_, a weighted rose plot was constructed using the angles and lengths of vectors representing individual neurons, after which a statistical test was performed in each angular bin. For analysis of the orthogonality of RDI values, the angles between the diagonal line and the vectors of fields with maximum RDI_SCN_ or RDI_CHC_ were obtained. Then, the product of their sine values, defined as the strength of RDI orthogonality, was obtained. This measurement was adopted because it had the characteristic that its value approached zero as any one of the fields came close to the diagonal line.

### Statistical analysis

Both the behavioral and neural data were analyzed using nonparametric statistical tests with the level of statistical significance set at α = 0.05 unless noted otherwise. Testing for statistical significance was two-sided, except when testing significance against a specific known value. For example, a one-sample Wilcoxon signed rank test was used to compare the behavioral performance for different scene stimuli against our performance criterion of 75% and to test whether the differences in RDI_SCN_ or RDI_CHC_ between field identification methods were significantly above zero. The proportional differences in cell types between two regions or two methods were tested using a Chi-square test. Comparisons of basic firing properties including mean firing rates, spatial information score and field width were conducted using a Wilcoxon rank sum test. The differences in slope and strength of theta phase precession were examined by 2-way mixed ANOVA with region as a between-subject and method as a within-subject factor, but an unpaired two-sample t test with Bonferroni correction was used for post hoc test because the number of observations for the within-subject factor was different across cells. Comparisons of ΔRDI_SCN_, ΔRDI_CHC_ and RDI orthogonality strength between the CA1 and subiculum were performed using the Wilcoxon rank sum test. In addition, the Wilcoxon rank sum test was also used for joint comparison of ΔRDI_SCN_ and ΔRDI_CHC_ in the vector-length-weighted rose plot to test regional differences in mean vector length within individual angular bins. Differences in RDI_SCN_ and RDI_CHC_ among cell types (i.e., MF ortho, MF non-ortho and SF) were assessed using a Kruskal-Wallis test, with the application of the Bonferroni-corrected Wilcoxon rank sum test for post hoc comparisons.

## Results

### Electrophysiological recording in the subiculum and CA1 in the VSM task

In the VSM task, rats (n = 5) learned to associate each scene stimulus with either a left or right turn response on the T-maze (**Figure 1A**). During recording sessions, rats performed the VSM task well above performance criterion (75%) for all stimuli (p-values < 0.0004 for all scenes, one-sample Wilcoxon signed-rank test; **Figure 1B**). Tetrodes located at the boundaries of either the CA1 or subiculum (including the border between them) were excluded from the analysis (**Figure 1C**). To quantify the anatomical distributions of recording locations for the CA1 and subiculum along the proximodistal axis, we measured the relative positions from which individual cells were recorded and normalized them across rats (**Figure 1D**). Only cells satisfying our unit-filtering criteria (CA1, n = 270; subiculum, n = 151) were used for analysis. Subicular cells were found along the entire proximodistal axis, whereas CA1 cells were mainly recorded from intermediate to proximal portions of the CA1. More details can be found in our previous study (Lee et al., 2018).

### Limitations of the firing rate-based method in detecting place fields in the subiculum

Prior studies (Barnes et al., 1990; Sharp and Green, 1994; Kim et al., 2012b; Lee et al., 2018) reported that cells in the subiculum fire at higher rates with lower spatial selectivity than those in the CA1, a finding also confirmed in our study. That is, cells recorded from the CA1 fired at focal and restricted locations along the T-maze (**Figure 2A**), whereas cells recorded from the subiculum tended to show broad and continuous firing fields (**Figure 2B**), making it challenging to identify a place field using the conventional field-detection method based on spatial firing rates. Specifically, although some subicular cells exhibited spatially tuned place fields similar to CA1 place fields (cells 234-4-1-5 and 232-5-4-8 in **Figure 2B**), some background spiking activity continued to occur outside their place fields. Furthermore, some subicular cells fired continuously across the entire track (cells 232-4-17-1 and 232-5-20-1 in **Figure 2B**), complicating efforts to define the field boundaries for these cells. These differences in field characteristics between the CA1 and subiculum can be more clearly observed in population rate maps constructed by stacking the rate maps of individual cells (**Figure 2A** and **2B**).

**Figure 2.**
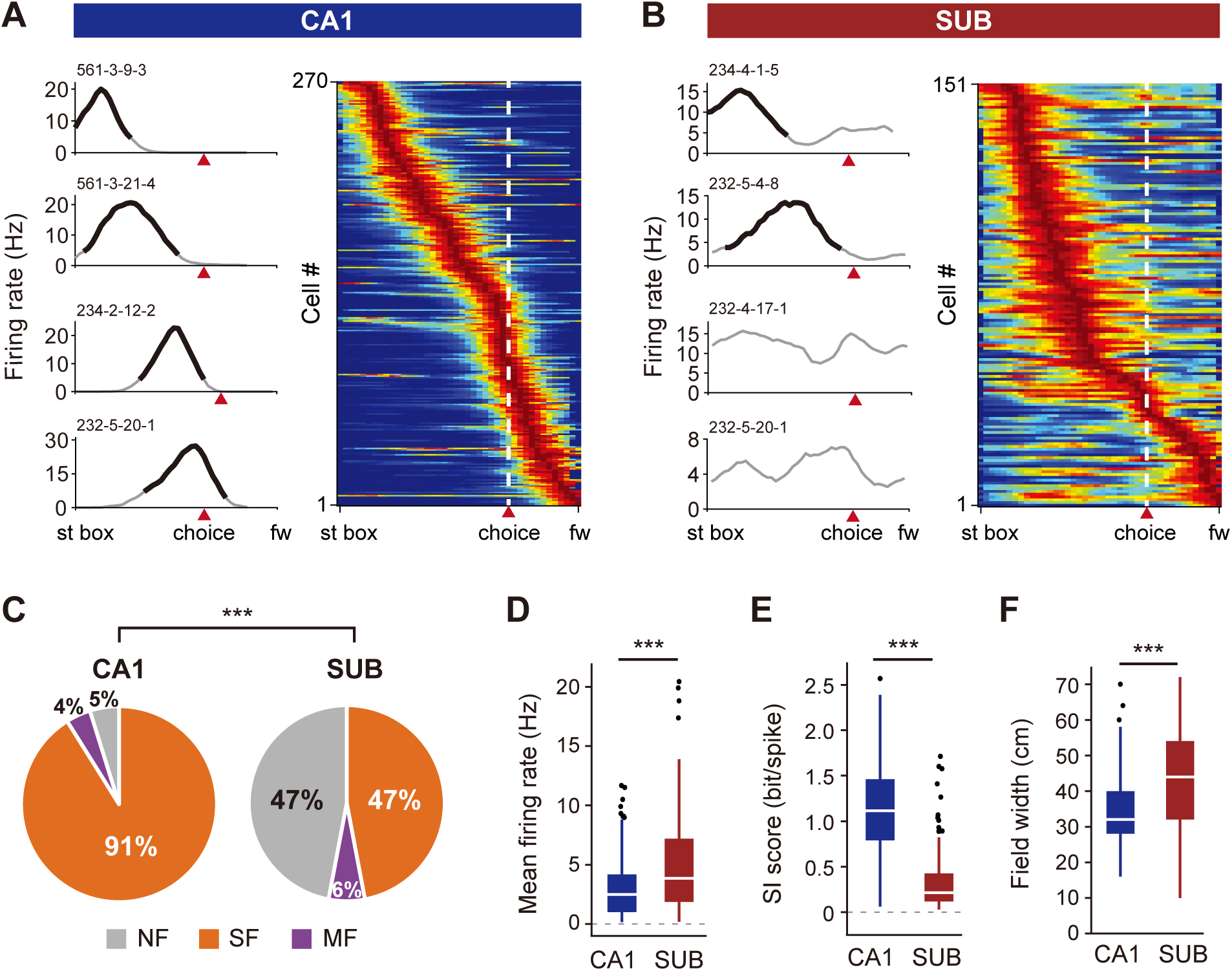
Poorer spatial firing patterns in the subiculum than the CA1. (**A**, **B**) Firing rate maps of single cells (left) and cell populations (right) in the CA1 (A) and subiculum (B), plotted as a function of linearized position on the T-maze from the start box (st box) to the food wells (fw) in both arms. Red arrowheads indicate the choice point after which rats’ positions diverged to the two arms. On the firing rate maps of single cells, verified place fields are overlaid with black lines, and non-place fields that did not pass the place field criteria are marked by black dotted lines. Serial numbers on the upper left corner are cell IDs. Indices on the right corner denote mean peak firing rates (Hz) and spatial information scores (bit/spike) of place and non-place fields. Population firing rate maps are sorted according to peak firing rate of each cell on the T-maze. White dashed lines and red arrowheads indicate the choice points. (**C**) Proportional differences of place cells defined by the firing rate-based method. Cells are classified into three groups according to the number of place fields within a cell: ‘single-field (SF)’ for one field, ‘multiple-field (MF)’ for more than one field, and ‘non-place field (NF)’ for zero field. ***p < 0.0001. (**D–F**) Differences in mean firing rate (D), spatial information score (SI score; E), and place field width (F) of recorded cells between the CA1 and subiculum. Box plot indicates interquartile range and median value for each region. ***p < 0.0001.

To quantitatively compare differential firing patterns between the two regions, we first classified cells according to the number of place fields: no place field, a single field, or multiple fields. A spatial firing distribution was considered a place field if its peak firing rate exceeded 1 Hz and its spatial information content (bits/spike) exceeded 0.5. Field boundaries were set at spatial bins in which the associated firing rates dropped below 33% of the peak firing rate (see **Methods**). Of cells that were active during the rat’s outbound journey on the T-maze, approximately 90% were single-field cells in the CA1, while only about half of cells exhibited either single- or multiple-field in the subiculum (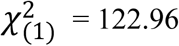, p < 0.0001; Chi-square test; **Figure 2C**). With respect to basic firing properties, cells in the CA1 showed lower firing rates (**Figure 2D**) with higher spatial information (**Figure 2E**) compared with those in the subiculum (firing rate, Z = 5.14, p < 0.0001; spatial information, Z = 14.2317, p < 0.0001; Wilcoxon rank-sum test). In addition, field width was larger in subicular place cells than in CA1 place cells (Z = 5.96, p < 0.0001; Wilcoxon rank-sum test) for both single-field and multiple-field cells (**Figure 2F**). Overall, these spatial firing pattern characteristics made it difficult to define place fields for individual neurons in the subiculum compared with those in the CA1.

### Identification of latent place fields based on theta phase precession of spiking

Our findings show that the fundamental differences in spatial firing characteristics between the CA1 and subiculum make it difficult to use conventional approaches commonly employed for analyzing place fields in both regions because these approaches have mostly been developed for place fields of cells in the hippocampus and not for those in the subiculum. In fact, a large number of subicular cells that would have been defined as no-field cells by conventional field-detection methods did fire more vigorously at particular locations of the track (**Figure 2B**, cells 232-4-17-1 and 232-5-20-1), but the conventional field-detection algorithm was unable to detect such spatial firing patterns because of the higher spontaneous firing activities throughout the track in subicular cells compared with CA1 neurons. Our previous study (Lee et al., 2018) tried to locate field boundaries in these cells by adjusting the threshold for detecting field boundaries or by finding local minima through statistical comparisons of trial-by-trial firing rates between neighboring bins. However, such methods still defined some subicular cells as having no fields. Furthermore, the conventional field-detection algorithm tended to ignore a small subfield if there was one dominant field with a very high firing peak.

To overcome such limitations, we explored the possibility of defining place fields using theta phase precession, a well-known phenomenon in which theta-related phases of spikes of a neuron gradually shift to earlier phases as the rat repeatedly passes through the cell’s place field (O’Keefe and Recce, 1993; Skaggs et al., 1996). In particular, we examined whether the broad firing field of a subicular neuron could be divided into multiple subfields if it were defined by theta phases of spikes. As shown in **Figure 3**, theta phase precession occurred robustly within the identified unitary place field in both the CA1 and subiculum as the rat ran along the track (CA1 single-field cells 234-2-12-2 and 561-2-3-1 in **Figure 3A**; subicular single-field cells 232-5-4-8 and 234-4-1-5 in **Figure 3B**). Importantly, those cells classified as having no place field exhibited multiple cycles of robust theta phase precessions (subicular non-place field cells 232-7-17-1 and 232-4-17-1 in **Figure 3C**).

**Figure 3.**
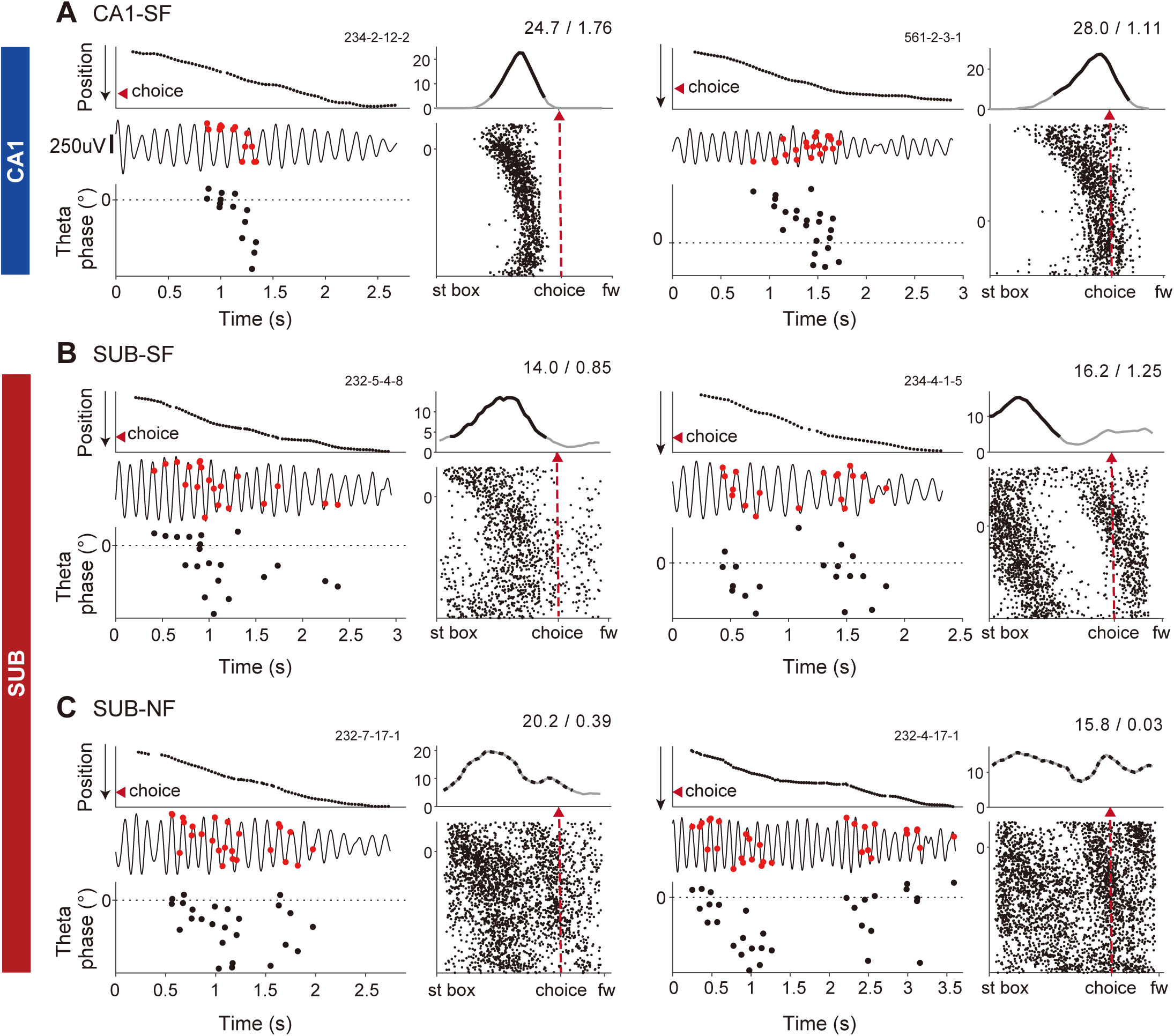
Robust and multiple theta phase precession in the subiculum. (**A-C**) Representative examples of theta phase precession in the CA1 single-field cells (A), subicular single-field cells (B), and subicular non-place field cells (C). The left column within each cell consists of linearized position (top), a raw trace of theta oscillation (middle), and spiking theta phases (bottom) in the temporal axis in a single trial. Spikes in the raw theta traces are marked by red circular dots. Scale bar, 250 μV. Spiking theta phases are plotted within a range of 360°, and the initial phase is adjusted for clear observation of theta phase precession. Serial numbers in the upper right corner are cell IDs. The right column displays a linearized firing rate map (top) and a position-phase plot (bottom) on the spatial axis across a session. Black solid lines overlaid on the firing rate maps indicate verified place fields, whereas black dotted lines are non-place fields. Numbers above firing rate maps denote peak firing rates (Hz) and spatial information scores (bit/spike) of place or non-place fields. Red arrowheads and red dashed lines mark choice points. Note that subicular cells showed multiple cycles of theta phase precession, some of which were as robust as those of CA1 cells.

To identify a spike cluster that belonged to a single theta-precession cycle in the phase-position plot, we used the DBSCAN (density-based spatial clustering with applications of noise) algorithm suggested by Ester et al. (1996) (see **Methods** for details). We compared the results of two different methods for detecting place fields: a firing rate-based method that finds a *rate-based field*, and the theta phase precession-based DBSCAN method, which finds a *phase-based field*. Both algorithms produced the same results in some cells in both the CA1 and subiculum (**Figure 4A**). However, we were also able to find new place fields for other cells based on the phase-based method. Specifically, some cells that were originally classified as single-field cells were converted into multiple-field cells by application of the phase-based algorithm (**Figure 4B** to **4D**). That is, in some cells, existing rate-based fields were subdivided into more than two phase-based fields (**Figure 4B**). In other cells, additional place fields that might not have been detectable by rate-based method (mostly owing to low firing peaks) were revealed by the phase-based clustering (**Figure 4C**). In a final group of cells, the phase-based method separated an existing field and simultaneously added a new field. (**Figure 4D**).

**Figure 4.**
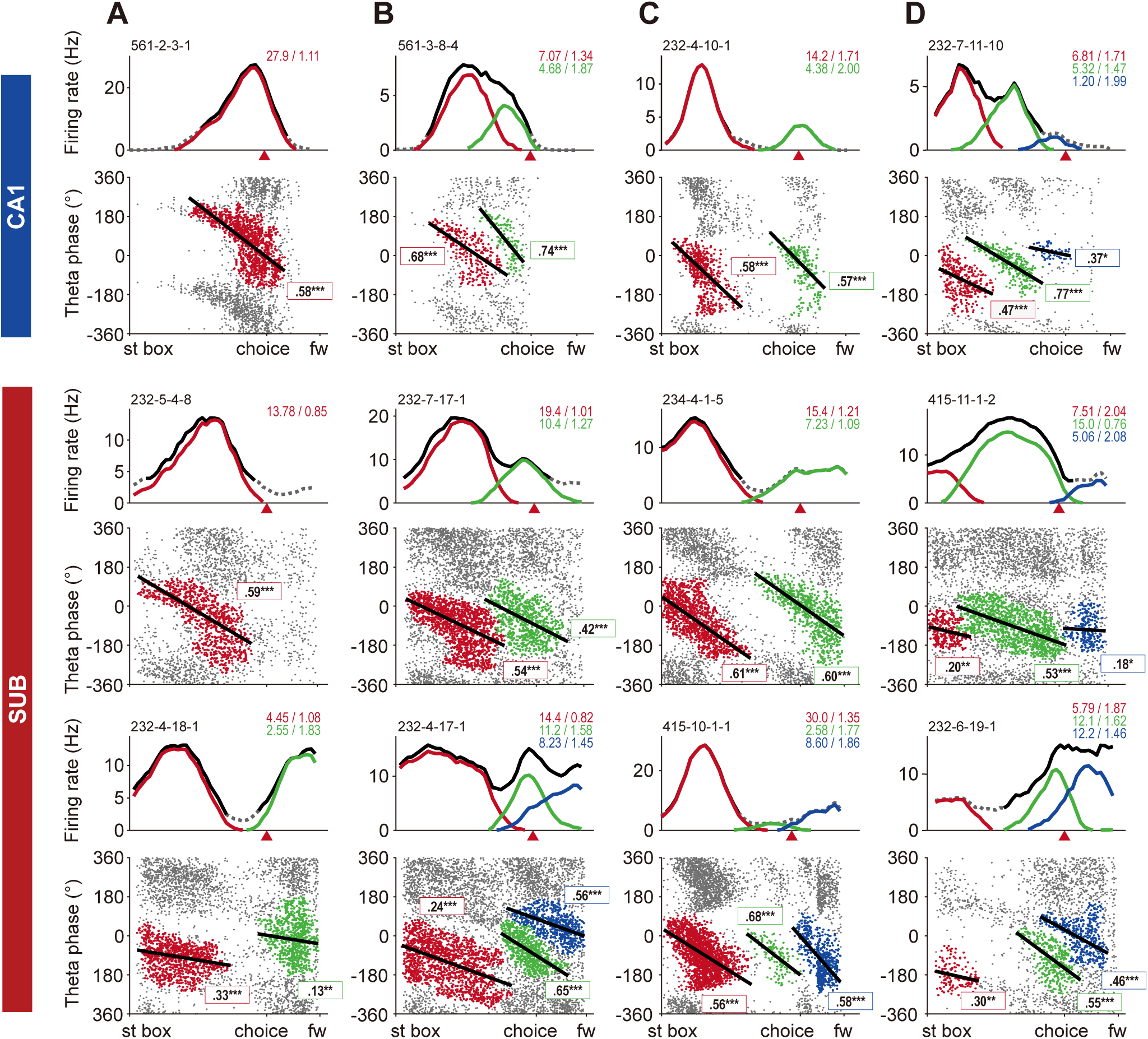
Identification of place fields based on spiking theta phases in the CA1 and subiculum. (**A–D**) Each subplot for individual cells consists of a linearized firing rate map (top) and a position-phase plot (bottom) on the spatial axis across a session. Gray lines on the firing rate maps indicate averaged firing activity, solid black lines denote place fields defined by firing rates (rate-based place fields), and color-coded lines denote place fields based on theta phases (phase-based place fields). Serial numbers above the firing rate maps are cell IDs. Color-coded numbers on the right corner indicate peak firing rate (Hz) and spatial information score (bit/spike) of individual phase-based place fields. Red arrowheads indicate choice points. Spike clusters in position-phase plots are color-coded with the same color used for the firing rate maps. Black straight lines on each spike cluster indicate the circular-linear regression line. The numbers and asterisks in the box with colored borders are circular-linear correlation coefficients; their significance for phase-based fields is indicated in the same color. ***p < 0.0001.

The proportions of cells showing different numbers of place fields changed using the phase-based clustering method compared with the rate-based method. Specifically, phase-based clustering classified 20% of CA1 cells and 62% of subicular cells as multiple-field cells and only 9% of subicular cells as having no fields (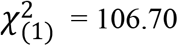, p < 0.0001; Chi-square test; **Figure 5A**). When the categorical changes were examined for each cell group, it turned out that three-quarters of the rate-based non-place cells in the CA1 and subiculum exhibited multiple phase-based place fields (**Figure 5B**). In addition, some rate-based single-field cells in the CA1 (14%) and subiculum (45%) were converted to multiple-field cells by the phase-based clustering. We also found that some rate-based multiple-field cells in the subiculum exhibited additional phase-based fields after applying the phase-based protocol (MF-added in **Figure 5B**). Phase-based place fields in the CA1 and subiculum still seemed to display some fundamental differences. For example, the widths of phase-based place fields remained significantly larger in the subiculum than in the CA1 (Z = 4.08, p < 0.0001; Wilcoxon rank-sum test; **Figure 5C**) compared with widths of rate-based fields.

**Figure 5.**
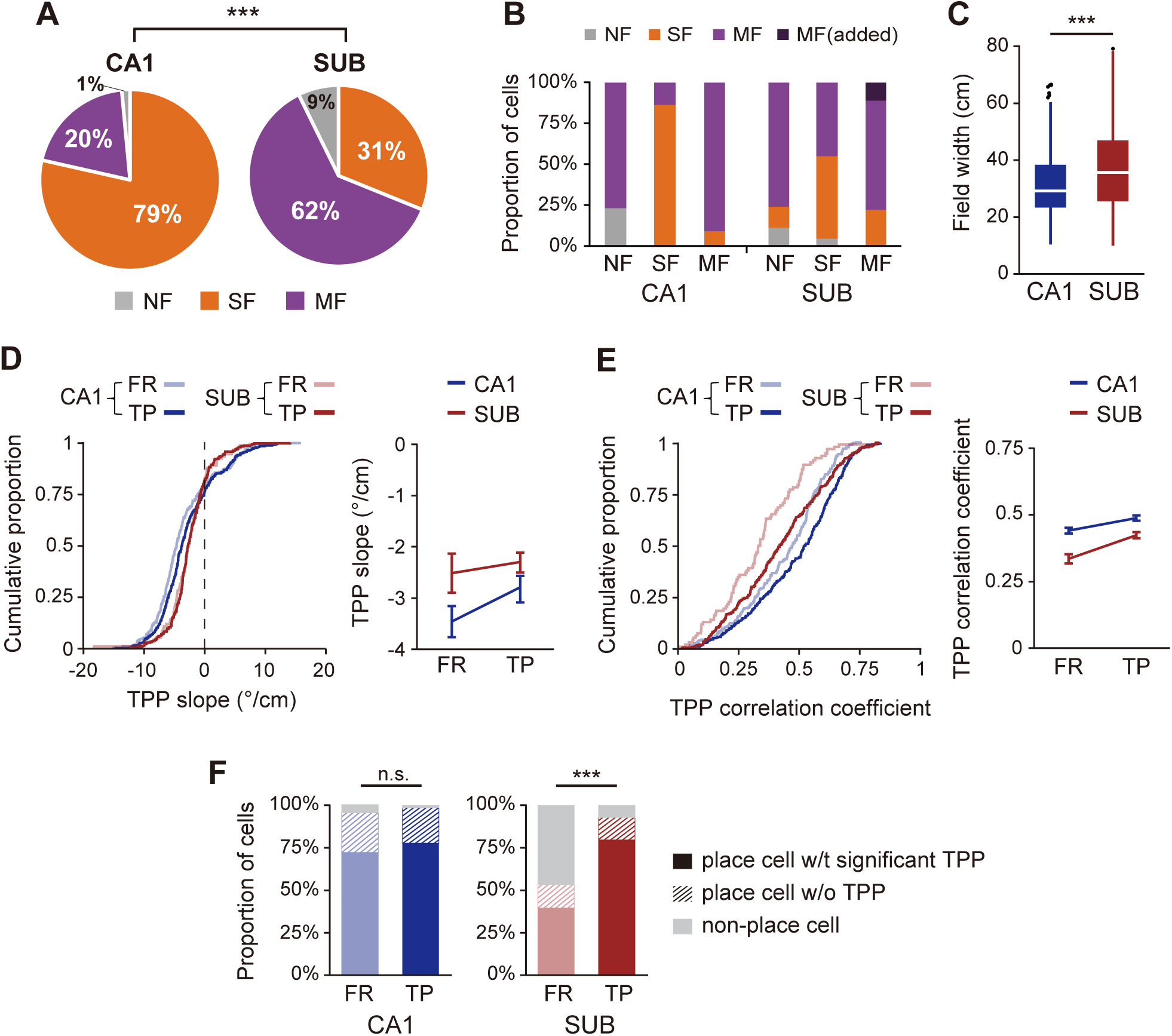
Advantages of spiking theta phase-based field identification. (**A**) Proportional differences of place cells defined using the theta phase-based method. Cells are classified into three groups according to the number of place fields within a cell as follows: ‘single-field (SF)’ for one field, ‘multiple-field (MF)’ for more than one field, and ‘non-place field (NF)’ for zero field. ***p < 0.0001. (**B**) Categorical changes of cells within each rate-based cell group (NF, SF, and MF on the *x*-axis) as the field identification method was shifted to phase-based. The bar graph shows what proportion of cells in each cell group classified by the rate-based method was re-categorized after the phase-based method. (**C**) Regional differences in place field width after phase-based field identification. ***p < 0.0001. (**D, E**) Cumulative distributions of theta phase precession (TPP) slope (D) and correlation coefficient (E) of place cells for each method (rate- and phase-based) and each region. Line graphs on the right side of each panel display mean values and standard errors for the same data. (**F**) Proportional changes in place cells with or without significant theta phase precession after the phase-based field identification within each region. ***p < 0.0001.

We next examined the robustness of theta phase precession of place cells in the subiculum compared with that in the CA1 using circular statistics (linear regression and linear correlation) for each spike cluster (Kempter et al., 2012). We found that the slope of theta phase precession was significantly different between the two regions (F_(1,435)_ = 4.43, p = 0.036), but it was not affected by the field-identification method (F_(1,613)_ = 0.59, p = 0.44, two-way mixed ANOVA with region as the between-subject factor and the field-identification method as the within subject factor; **Figure 5D**). There was no interaction between the region and field-detection method (F_(1,613)_ = 0.52, p = 0.47), mostly attributable to the reduced regional difference when the phase-precession slope was calculated based on the phase-based fields compared to the rate-based fields. The precession slope of rate-based fields tended to be steeper in the CA1 than in the subiculum (t_(805)_ = 2.01, p = 0.045 for Bonferroni-corrected unpaired two-sample t-test; corrected α = 0.0125), an outcome that could be expected based on the larger field width of subicular place cells. However, the regional difference in slope diminished when using the phase-based method (t_(496)_ = 1.64, p = 0.102). The slope of theta phase precession was not affected by the field-detection method within each region (CA1, t_(555)_ = 1.39, p =0.16; SUB, t_(637)_ = 0.03, p =0.98). On the other hand, the strength of theta phase precession of place cells evaluated by circular-linear correlation coefficient was significantly different between the CA1 and subiculum (F_(1,445)_ = 31.57, p < 0.0001) and between the two field-detection methods (F_(1,655)_ = 50.25, p < 0.0001; two-way mixed ANOVA; **Figure 5E)**. The interaction between the region and method was not significant (F_(1,655)_ = 3.68, p = 0.055). Post-hoc tests revealed that the phase-precession strength increased in phase-based fields compared with rate-based fields in both regions (CA1, t_(565)_ = 4.77, p < 0.0001; subiculum, t_(695)_ = 5.36, p < 0.0001; unpaired two-sample t test with Bonferroni correction; corrected α = 0.0125). Although precession strength was significantly lower in the subiculum than in the CA1 even after phase-based field identification (rate-based, t_(896)_ = 5.08, p < 0.0001; phase-based, t_(541)_ = 4.03, p < 0.0001), the precession strength in the subiculum increased closer to that of the CA1.

We also tested whether the theta phase-based parsing of subicular broad firing fields changed the proportion of place cells with significant theta phase precession. A place cell was classified as having significant theta phase precession as one of its subfields met the following criteria: range of phase shift is equal to or larger than 180°; slope of circular-linear regression line is lower than zero; and p-value of circular-linear correlation is equal to or lower than 0.05. Then, we obtained the proportion of place cells with or without significant theta phase precession and even non-place cells. The proportional changes between field identification methods were significant in the subiculum (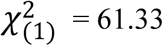, p < 0.0001; Chi-square test; **Figure 5F**), but not in the CA1 (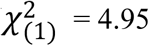, p = 0.084). Taken together, based on the theta phase precession cycle, these findings indicate that the DBSCAN algorithm effectively identified the multiple subfields enveloped in the broad firing activities of the subicular cells.

### Increase in task-relevant information in phase-based fields of subicular neurons

We previously reported that firing of neurons in the CA1 and subiculum was correlated with the visual scene stimulus and choice response in the VSM task in the form of rate remapping (Delcasso et al., 2014; Lee et al., 2018). Here, we examined whether scene- or choice-dependent rate remapping also appeared in phase-based fields of neurons in the CA1 (n = 211) and subiculum (n = 139). To quantify rate remapping, we obtained a rate difference index (RDI) for individual rate-based and phase-based fields using the firing rate maps associated with different task conditions (see **Methods**; **Figure 6A**). The RDI for choice response (RDI_CHC_) was calculated using only the spiking activity recorded up to the choice point. In contrast, for the scene-based RDI (RDI_SCN_), spiking activity associated with places beyond the choice point were also included because only the scenes associated with the same choice arm were compared. Place fields representing only arm areas were excluded from the RDI analysis.

**Figure 6.**
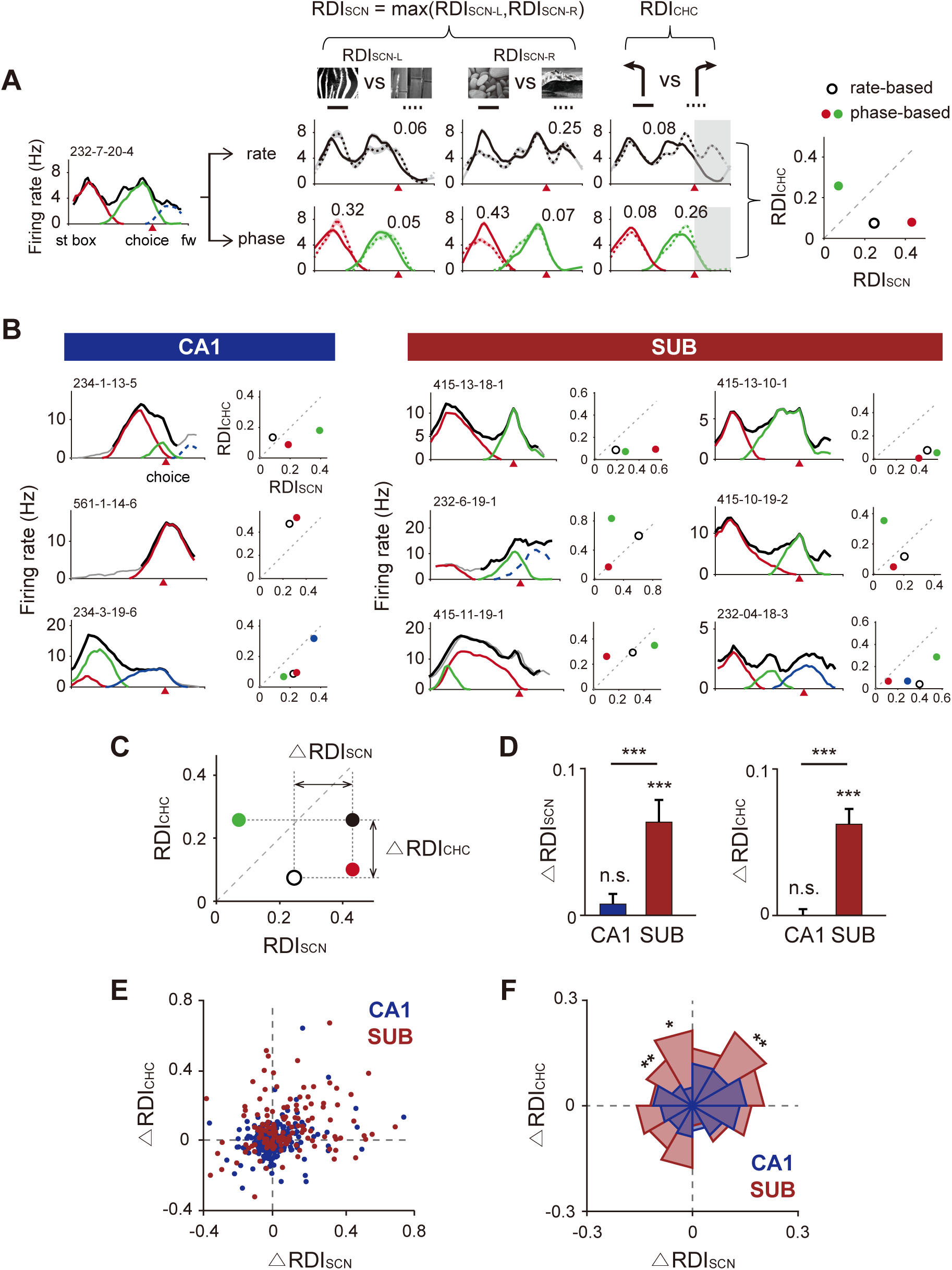
Scene- and choice-dependent rate remapping is enhanced in the subiculum but not in the CA1 after phase-based field identification. (**A**) A representative subicular cell illustrating how difference indices for scene (RDI_SCN_) and choice (RDI_CHC_) are obtained. The linearized firing rate map in the left panel shows rate-based fields (black line) and phase-based fields (color-coded lines) averaged across all trials. Middle panel shows firing rate maps associated with different task-relevant information. Rate-based fields are marked as black lines (upper row) and phase-based fields are color-coded (bottom row). Shaded areas overlaid on the fields are standard errors. Numbers above the fields indicate RDI values. Rightmost panel shows RDI_SCN_ and RDI_CHC_ values for individual fields marked as dots on the scatter plot; open black dots correspond to rate-based fields and color-coded dots denote phase-based fields. The dashed line shows the diagonal. (**B**) Example neurons in the CA1 and subiculum with their RDI values associated with the scene and choice information. Within each neuron, linearized firing rate map (left) and RDI scatter plot (right) are shown as in (A). Solid black lines on firing rate maps are firing rate (FR)-based firing fields and color-coded lines are theta phase (TP)-based place fields, with arm fields depicted in dotted lines. Serial numbers above the rate maps are cell IDs. Red arrowheads indicate choice points. (**C**) Illustration showing how RDI differences (i.e., ΔRDI_SCN_ and ΔRDI_CHC_) are measured using the rate-based method (open black circle) and phase-based method (closed black circle). The closed black circle is a representative point for the phase-based method, marked by selecting maximum values among RDIs obtained from all phase-based fields. (**D**) Bar graphs comparing the magnitude of changes in RDI_SCN_ and RDI_CHC_ between regions. Data are presented as means ± standard error of the mean. ***p < 0.0001. (**E**) Scatter plot jointly displaying ΔRDI_SCN_ and ΔRDI_CHC_ for all neurons in the CA1 and subiculum. Note that subicular neurons are more dispersed in the first and second quadrant than CA1 neurons. (**F**) Weighted rose plot constructed using data from (E) for statistical comparison. **p < 0.01, *p < 0.05.

We found that the phase-based field identification method extracted task-relevant information more clearly than the rate-based method, especially in the subiculum. It also revealed new information that went undetected by the conventional rate-based method. For example, the phase-based method identified two fields for a subicular cell shown in **Figure 6A** on the stem of the maze that were unidentifiable by the conventional rate-based method. One of the phase-based fields (red in **Figure 6A**) showed a larger amount of scene information than the rate-based field (0.32 > 0.06 for RDI_SCN-L_ and 0.43 > 0.25 for RDI_SCN-R_). The other phase-based field (green in **Figure 6A**) showed minimal information on the visual scene but carried more information on the choice response compared with the rate-based field (0.26 > 0.08 for RDI_CHC_). As illustrated by the neuronal examples in **Figure 6B**, some phase-based fields showed stronger rate remapping for scenes than for choices (cell 234-1-13-5 in CA1, cells 415-13-18-1, 415-13-10-1 in the subiculum), whereas other phase-based fields exhibited the opposite pattern (cell 561-1-14-6 in CA1; cells 232-6-19-1 and 415-10-19-2 in the subiculum). Furthermore, scene and choice information increased to a similar degree in some phase-based fields (**Figure 6B**, cell 234-3-19-6 in CA1; cells 415-11-19-1 and 232-4-18-3 in the subiculum).

We next investigated the extent to which task-related information carried by a single unit changed when the field identification protocol was changed from the rate-based to the phase-based method. For this purpose, if one cell showed multiple fields, the maximum RDI value was selected as the representative RDI of the cell (closed black dot in **Figure 6C**). Then, we calculated the *difference in RDI* (ΔRDI) by subtracting the representative RDI value of the rate-based protocol from the representative RDI value of the phase-based protocol for scene (ΔRDI_SCN_) and choice (ΔRDI_CHC_) information, respectively. Both RDI_SCN_ and RDI_CHC_ increased remarkably in the subiculum after theta phase-based field identification (T = 1582, p = 0.0002 for ΔRDI_SCN_; T = 2415, p < 0.0001 for ΔRDI_CHC_; one-sample Wilcoxon signed rank test), but no significant increase was found in the CA1 (T = 365, p = 0.68 for ΔRDI_SCN_; T = 645, p = 0.47 for ΔRDI_CHC_; **Figure 6D**). The RDI increases for subicular neurons were significantly higher than those for CA1 neurons for both visual scenes (Z = 3.26, p = 0.0011 for ΔRDI_SCN_) and choices (Z = 5.44, p < 0.0001 for ΔRDI_CHC_; Wilcoxon rank-sum test).

For joint comparisons of changes in scene- and choice-based rate remapping, the differences in RDI_SCN_ and RDI_CHC_ of each cell were marked as dots on a scatter plot (**Figure 6E**). As shown in the first and second quadrants of the scatter plot, RDI_SCN_ and RDI_CHC_ values increased jointly after the phase-based field identification method in the subiculum compared with the CA1, where there was no such trend. Furthermore, RDI_CHC_ values were enhanced in some cells irrespective of RDI_SCN_ values. To statistically compare the regional differences visible in the scatter plot, we first divided the quadrant into equally spaced angular bins (30°) and drew the mean vectors for each bin in a vector-length–weighted rose plot. As shown in **Figure 6F**, the stronger RDI enhancement in the subiculum compared with the CA1 was clearly visible in the first and second quadrants (bin 30°-60°, p = 0.006; bin 90°-120°, p = 0.014; bin 120°-150°, p = 0.005; Wilcoxon rank-sum test). Taken together, these results indicate that the theta phase-based field detection method is capable of identifying task-relevant information that would otherwise have been unidentifiable using the traditional rate-based field-detection protocol.

### Neurons in the subiculum represent scene and choice information more orthogonally in their multiple phased-based fields than CA1 cells

We further examined the functional significance of amplified task-related information (i.e., scene and choice) discovered by the phase-based method in subicular neurons compared with CA1 cells in our VSM task. If a place field showed the same amount of rate modulation for both scene and choice factors, the corresponding data point on the RDI scatter plot should be located on the diagonal (e.g., field 1 in **Figure 7A**). In contrast, if the amount of field remapping was influenced to a greater degree by one of the task-related factors, the data point should be located farther away from the diagonal (e.g., field 2 in **Figure 7A**). If a cell had multiple phase-based fields and each field represented either scene or choice information more strongly than the other, the cell was considered as *orthogonalizing* scene and choice information (**Figure 7B**).

**Figure 7.**
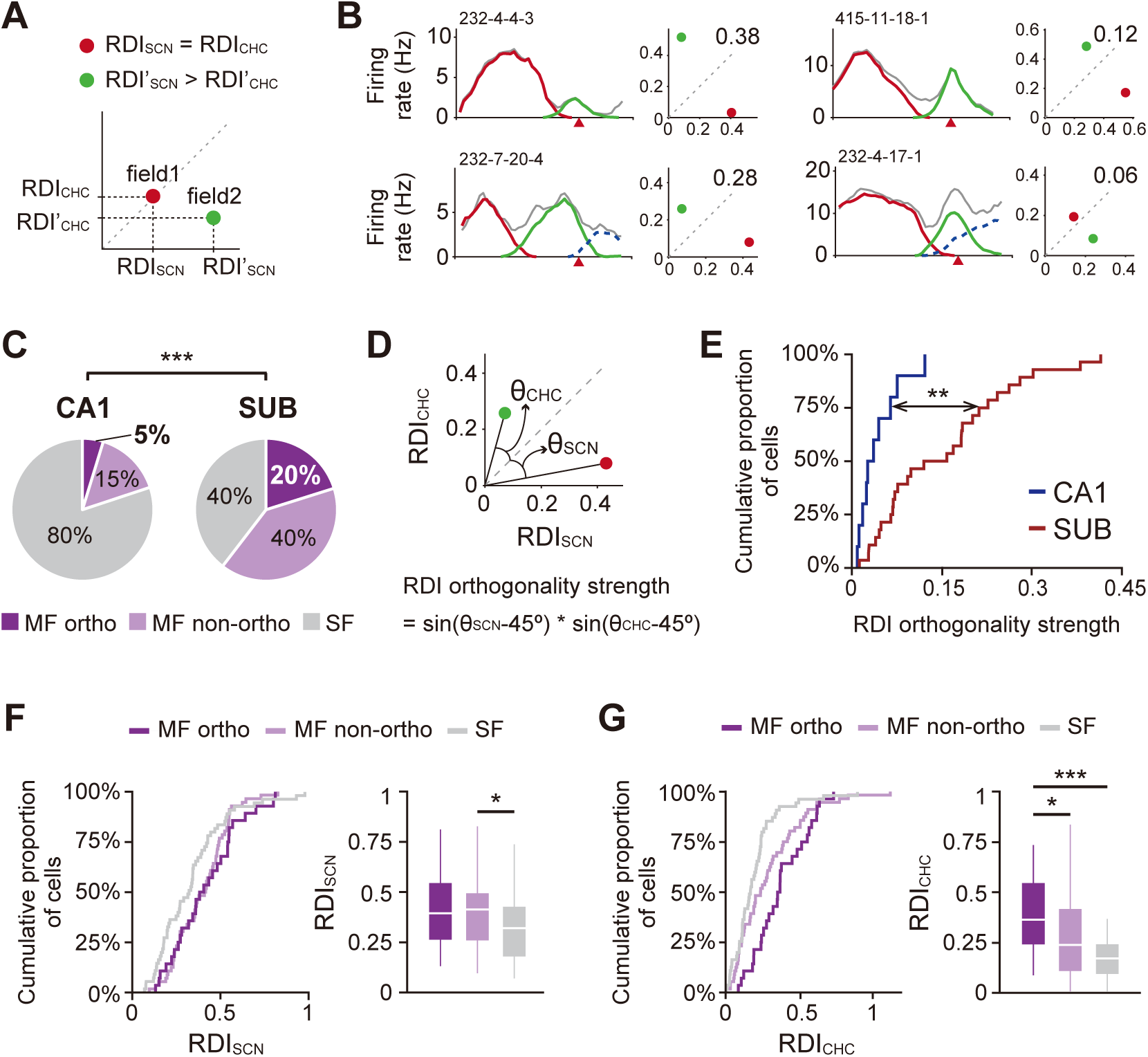
Scene and choice information are orthogonally represented by multiple phase-based fields of subicular neurons. (**A**) Illustration showing the different relationships between RDI_SCN_ and RDI_CHC_ of example fields on the RDI scatter plot. Field1 (red) near the diagonal line shows the same amount of rate modulation for scene and choice information, whereas field2 (green) located further away from the diagonal line had much stronger rate-remapping for scene than choice information. (**B**) Four examples of orthogonal representations for scene and choice information for individual neurons. For each neuron, the left panel shows a linearized firing rate map (left) and the right panel shows an RDI scatter plot. Each phase-based field is color-coded. Serial numbers above the rate map indicate cell IDs. Numbers on the scatter plots indicate RDI orthogonality strength. (**C**) Proportion of cells for which phase-based fields have orthogonal representations for scene and choice information. ***p < 0.0001. (**D**) Illustration displaying how the strength of orthogonal representations for scene and choice information is quantified. θ_SCN_ and θ_CHC_ indicate the angles between the diagonal line and the vectors of the fields whose RDI_SCN_ or RDI_CHC_ is the maximum value. (**E**) Cumulative distribution of RDI orthogonality strength for each region. **p < 0.01. (**F**, **G**) Cumulative proportion of subicular cells for RDI_SCN_ (F) and RDI_CHC_ (G). Bar graphs on the right side of each panel show RDI differences between subgroups within the subiculum. ***p < 0.0001, *p < 0.016.

A significantly larger portion of subicular neurons exhibited orthogonal representations of task-related variables than CA1 neurons (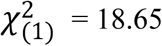, p < 0.0001; Chi-square test; **Figure 7C**). To test whether the orthogonal representation occurred more strongly in subicular cells, we measured the angle between the diagonal and the vector of each phase-based field (θ_SCN_ and θ_CHC_; **Figure 7D**), then calculated orthogonality strength by multiplying the sine values of the angles. RDI orthogonality was significantly stronger in the subiculum than in the CA1 (Z = 3.3, p = 0.001, Wilcoxon rank-sum test; **Figure 7E**).

Finally, we tested whether the amount of rate remapping differed between the following cell groups: multiple-field cells with orthogonal representations (MF ortho), multiple-field cells without such representations (MF non-ortho), and single-field cells (SF). Since the number of CA1 cells showing orthogonal representations was too small (n = 10) to obtain sufficient statistical power, tests were performed only within the subiculum. There were significant differences in both scene (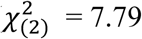, p = 0.02, Kruskal-Wallis test; **Figure 7F**) and choice (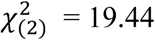, p < 0.0001; **Figure 7G**) between the subgroups. Specifically, cells having multiple fields showed larger RDI_SCN_ values than those with single fields (MF ortho vs. SF: Z = 2.12, p = 0.034; MF non-ortho vs. SF: Z = 2.5, p = 0.013; MF ortho vs. MF non-ortho: Z = 0.37, p = 0.7; Wilcoxon rank-sum test with Bonferroni correction; corrected *α* = 0.016). Moreover, subicular cells with orthogonal representations exhibited significantly larger RDI_CHC_ values than other groups (MF ortho vs. SF: Z = 4.51, p < 0.0001; MF non-ortho vs. SF: Z = 2.16, p = 0.031; MF ortho vs. MF non-ortho: Z = 2.4, p = 0.016). These findings indicate that subicular multiple fields identified based on the relationships between spiking theta phase precessions did not uniformly represent task-relevant information. Instead, they carried heterogeneous task-related information in a more orthogonal fashion compared with CA1 cells. Furthermore, such subicular cells represented task-related information more strongly than CA1 cells in the VSM task.

## Discussion

In the current study, we characterized the firing patterns of place cells in the CA1 and subiculum using both phase- and rate-coding methods. Our findings demonstrate that some of the broad place fields of subicular neurons can be parsed into multiple fields using the theta-phase precession cycle. The newly discovered, phase-based place fields in the subiculum were more similar to those in CA1 in terms of field size and phase-precession strength. However, unlike the case in the CA1, the neural representational strength of task-relevant information was significantly improved in the subiculum by the phase-based field-detection method. Furthermore, our results suggest that firing for multiple fields by a single neuron may provide the subiculum with the unique function of representing different types of task-related information in an orthogonal fashion compared with the CA1.

### Underlying mechanisms of multiple cycles of theta phase precession and their associated place fields in the subiculum

One possible mechanism underlying the multiple cycles of theta phase precession and their associated place fields in the subiculum is convergent inputs from multiple place cells in the CA1 to a subicular cell. To our knowledge, whether a single subicular neuron is innervated by multiple CA1 place cells is still largely unknown. However, it has been reported that axon branches extending from a single CA1 pyramidal cell diverge to a very wide region within the subiculum, covering approximately 2 mm along the septotemporal axis (Tamamaki et al., 1987) and one-third of the subiculum along the proximodistal axis (Amaral et al., 1991). In addition, approximately 40% of CA1 pyramidal cells are known to send efferent projections to the subiculum (Roy et al., 2017). Based on these anatomical results, it is possible that a single subicular cell receives synaptic inputs from multiple CA1 pyramidal cells. If this is the case, a subicular place cell that receives inputs from multiple place cells in the CA1 whose firing peaks are located at distant locations may develop multiple place fields. Conversely, if multiple CA1 place cells sending projections to a single subicular place cell have overlapping place fields, then the subicular cell might exhibit a single broad firing field. Some prior studies may support these possibilities (Fernandez-Ruiz et al., 2017; Jaramillo and Kempter, 2017).

Another possibility is that the multiple fields of the subiculum might be based on inputs from cells in the medial entorhinal cortex, especially grid cells showing periodic firing fields and theta phase precession. Some models have shown that theta phase precession in the CA1 could be derived from grid cells in the medial entorhinal cortex (Molter and Yamaguchi, 2008; Schlesiger et al., 2015). It has also been reported that temporal coding (including theta phase precession) in the CA1 is impaired by lesioning of the medial entorhinal cortex (Schlesiger et al., 2015). However, cells in layer 3 of the entorhinal cortex, which mainly project to the CA1 and subiculum, do not exhibit phase precession relative to theta rhythm (Hafting et al., 2008). Whether theta phase precession in the subiculum is inherited from grid cells in the medial entorhinal cortex remains to be investigated.

Lastly, there is the possibility that cells in the subiculum might be influenced by multiple sources of theta rhythm—one from an extracellular source and another generated intrinsically. Specifically, some previous studies have proposed an interference model as the mechanism for theta phase precession in the CA1 (O’Keefe and Recce, 1993; Burgess et al., 1994; O’Keefe and Burgess, 2005). According to this model, there is an intrinsic theta oscillator within pyramidal cells that causes theta phase precession while maintaining a frequency that can be different from that of the extracellular theta rhythm. This model is supported by experimental evidence showing that pyramidal cells in the dorsal hippocampus show higher intrinsic oscillation frequencies than those in the ventral hippocampus, resulting in smaller place fields in the dorsal hippocampus (Maurer et al., 2005). Experimental evidence for the presence of an intrinsic theta oscillator in the subiculum has not been reported. However, because principal cells in the subiculum have denser recurrent connectivity than those in the CA1 (Harris and Stewart, 2001; Harris et al., 2001; Bohm et al., 2015), it is possible that cells in the subiculum can generate local rhythms intrinsically. Notably, a recent study reported that an atypical type of sharp-wave ripple occurs in the subiculum, independent of traditionally known CA3-originating sharp-wave ripples (Imbrosci et al., 2021), an observation that may support the possibility that subicular neurons intrinsically generate their own local oscillations.

### Clustering algorithm for identifying multiple cycles of theta phase precession and their associated place fields

A previous study by Maurer et al. (2006) demonstrated that the partially overlapping place fields of a single cell in the CA1 could be segmented by manually drawing boundaries around the spikes belonging to individual cycles of theta phase precession on the position-phase scatter plot. Further improving this strategy, Kim et al. (2012b) developed an automated algorithm that constructs a phase-position firing-rate map from normalized phase-position plots of rat occupancy and then defines place fields based on detection of local maxima. However, there were challenges to adopting this previous protocol in the current study. First, these authors used a behavioral paradigm in which rats ran along a track in the absence of environmental change or mnemonic task demand, whereas in the current study rats performed a mnemonic task in which they were required to associate different scenes with discrete behavioral choices. Our previous study showed that firing rates of subicular cells are modulated in relation to task-related information (i.e., scene and choice) (Lee et al., 2018). In that case, even if a field had a high firing rate in one condition, it might show a low firing rate in the other condition. Accordingly, in the overall firing rate map obtained by averaging the firing rates for all conditions, the high firing rate under one specific condition is likely to be canceled out by that under the other condition, making it difficult to definitively establish peak firing and field boundaries when defining a place field. Second, the length of the linear track used in the current study was relatively short, possibly resulting in more overlap between place fields and thus creating difficulties in setting an appropriate threshold for segmenting individual fields on the phase-position firing-rate map.

Therefore, to eliminate the risk of being unable to find potential task-related firing fields, we adopted the clustering algorithm DBSCAN (Ester et al., 1996), which can be applied to the raw phase-position spiking plot without normalizing for occupancy. We chose this algorithm for several reasons. First, when sampled sufficiently, spikes tend to occur at the most preferred location within a place field with highest probability and then gradually diminish as the distance from the field center increases (i.e., Gaussian-like distribution).

Because of its density-based algorithmic nature, the DBSCAN algorithm is suitable for finding clusters when data points exhibit such distributions. Second, DBSCAN has the advantage of robustly detecting outliers, which enabled us to process continuous and spontaneous firing activities of subicular fields. Furthermore, DBSCAN does not require an experimenter to predetermine the number of clusters. Finally, the DBSCAN algorithm does not limit cluster shape, so it can flexibly find clusters in a complex data set.

### Functional significance of the more orthogonal representation of scene and choice information in the subiculum than the CA1

The spatial firing properties of subicular neurons are different from those of CA1 cells. This unique nature of the subiculum has been attributed to signals from outside the hippocampal formation, including those related to movement and head direction of the thalamic region (Frost et al., 2021). However, task-related information such as scene and choice information in the subiculum should also be influenced by inputs from the dorsal CA1, as demonstrated in our previous studies (Kim et al., 2012a; Delcasso et al., 2014). This possibility is supported by findings of the current study showing significantly enhanced rate modulation for task-relevant information, as only the group of spikes constituting a single cycle of theta phase precession was extracted for measuring the representational strength of task-related information.

The enhancement of task-related signals in the subiculum allowed us to investigate the functional roles of subicular cells in the VSM task compared with our previous study in which we relied on the traditional rate-based field-detection algorithm (Lee et al., 2018). Remarkably, some subicular neurons carried orthogonal scene and choice information in their subfields. This phenomenon could arise in a scenario in which different CA1 cells, each carrying one type of task-related information more strongly than the other, send their outputs to a single subicular neuron. Subicular neuron could then facilitate associative learning by representing different types of information concurrently so that downstream structures receive more associative information between the critical task variables. Note that a subicular cell tends to represent different task-related information in separated fields associated with distant locations, but not conjunctively in one field. The different types of information represented by the separate fields of a subicular cell might then be transmitted into identical target regions nearly simultaneously with a certain phase relationship, potentially contributing to the formation and retention of hippocampal associative memory.

### Functional subclasses of neurons in the subiculum may play key roles in hippocampal-dependent action in a visual contextual memory task

Numerous studies have suggested the presence of anatomical and physiological subpopulations in the subiculum. Specifically, it has been reported that afferent and efferent projections of the subiculum are organized topographically along the proximodistal axis (Tamamaki et al., 1987; Amaral et al., 1991; Tamamaki and Nojyo, 1995; Naber and Witter, 1998; Witter et al., 2000; Ishizuka, 2001; Witter, 2006; Cembrowski et al., 2018b; Kitanishi et al., 2021), as the proximal and distal parts of the subiculum are clearly divided according to gene expression in principal cells (Cembrowski et al., 2018b; Cembrowski et al., 2018a). In addition to these anatomical subdivisions, *in vitro* physiological studies have reported two types of intrinsic firing for subicular principal cells—bursting and regular-spiking (Stewart and Wong, 1993; Taube, 1993; Behr et al., 1996)—and shown that these cells exhibit a unique distribution gradient along the proximodistal axis (Harris et al., 2001; Jarsky et al., 2008; Kim and Spruston, 2012). These two classes of cells are modulated differently by sharp-wave ripples of the CA1 and have different intrinsic connectivity (Bohm et al., 2015). Furthermore, a series of recent *in vivo* studies identified subpopulations in the subiculum with different spatial firing characteristics (Brotons-Mas et al., 2017; Roy et al., 2017; Simonnet and Brecht, 2019; Poulter et al., 2021).

Our physiological results also suggest that there are functionally different subclasses of neurons within the subiculum. In our previous paper, subicular cells with a single broad field showed a “schematic” firing pattern that depended on the cognitive structure of the task; we speculated that this firing pattern serves to mediate contextual behavior by representing the discrete region associated with critical epochs of the task (Lee et al., 2018). That is, cells in the CA1 and subiculum represent specific location information and an epoch-based region, respectively. On the other hand, subicular cells with multiple focal fields are thought to contribute to associative memory by subsequently—but almost concurrently—transmitting different types of task-relevant information to downstream structures where choice-related actions and decisions occur.

Taken together, our findings indicate that the subiculum may support visual contextual behavior in space through two processes, each driven by a distinctive neuronal class. Information regarding context and future path leading to the goal location, separately recognized at focal and distant place fields of the CA1, could be integrated by subicular cells with multiple phase-based fields and transmitted to downstream areas. In particular, such information processing in the subiculum could be critical for converting contextual information into an action signal. At the same time, subicular cells with a broad single field may represent the area in which all locations are associated with a common task-related variable, such as a specific visual scene or choice response. A recent study showing that CA1-projecting subicular cells receive direct inputs from the visual cortex and send their projections to some critical regions (e.g., perirhinal cortex, CA1) in the VSM task used in the current study may also support the functional significance of the subiculum in the visual contextual behavioral task (Sun et al., 2019).

## Acknowledgments

This study was supported by the National Research Foundation of Korea (NRF 2017M3C7A1029661, 2018R1A4A1025616, and 2019R1A2C2088799) and the BK21, FOUR by the Ministry of Education and the NRF.

